# Beyond RuBisCO: Convergent molecular evolution of multiple chloroplast genes in C_4_ plants

**DOI:** 10.1101/2021.08.30.457919

**Authors:** Claudio Casola, Jingjia Li

## Abstract

**Background:** The recurrent evolution of the C_4_ photosynthetic pathway in angiosperms represents one of the most extraordinary examples of convergent evolution of a complex trait. Comparative genomic analyses have unveiled some of the molecular changes associated with the C_4_ pathway. For instance, several key enzymes involved in the transition from C_3_ to C_4_ photosynthesis have been found to share convergent amino acid replacements along C_4_ lineages. However, the extent of convergent replacements potentially associated with the emergence of C_4_ plants remains to be fully assessed. Here, we introduced a robust empirical approach to test molecular convergence along a phylogeny including multiple C_3_ and C_4_ taxa. By analyzing proteins encoded by chloroplast genes, we tested if convergent replacements occurred more frequently than expected in C_4_ lineages compared to C_3_ lineages. Furthermore, we sought to determine if convergent evolution occurred in multiple chloroplast proteins beside the well-known case of the large RuBisCO subunit encoded by the chloroplast gene *rbcL*.

**Methods:** Our study was based on the comparative analysis of 43 C_4_ and 21 C_3_ grass species belonging to the PACMAD clade, a focal taxonomic group in many investigations of C_4_ evolution. We first used protein sequences of 67 orthologous chloroplast genes to build an accurate phylogeny of these species. Then, we inferred amino acid replacements along 13 C_4_ lineages and 9 C_3_ lineages using reconstructed protein sequences of their ancestral branches, corresponding to the most recent common ancestor of each lineage. Pairwise comparisons between ancestral branches allowed us to identify both convergent and divergent amino acid replacements between C_4_-C_4_, C_3_-C_3_ and C_3_-C_4_ lineages.

**Results:** The reconstructed phylogenetic tree of 64 PACMAD grasses was characterized by strong supports in all nodes used for analyses of convergence. We identified 217 convergent replacements and 201 divergent replacements in 45/67 chloroplast proteins in both C_4_ and C_3_ ancestral branches. Pairs of C_4_-C_4_ ancestral branches showed higher levels of convergent replacements than C_3_-C_3_ and C_3_-C_4_ pairs. Furthermore, we found that more proteins shared unique convergent replacements in C_4_ lineages, with both RbcL and RpoC1 (the RNA polymerase beta’ subunit 1) showing a significantly higher convergent/divergent replacements ratio in C_4_ branches. Notably, significantly more C_4_-C_4_ pairs of ancestral branches showed higher numbers of convergent vs. divergent replacements than C_3_-C_3_ and C_3_-C_4_ pairs. Our results demonstrated that, in the PACMAD clade, C_4_ grasses experienced higher levels of molecular convergence than C_3_ species across multiple chloroplast genes. These findings have important implications for both our understanding of the evolution of photosynthesis and the goal of engineering improved crop varieties that integrates components of the C_4_ pathway.

## Introduction

Convergent evolution represents the independent acquisition of similar phenotypic traits in phylogenetically distant organisms. Understanding the genomic changes underlying the recurrent emergence of phenotypes is a major goal of molecular evolution. The rapidly increasing taxonomic breadth of genomic resources combined with the development of rigorous frameworks to comparatively investigate molecular changes has accelerated the pace of discovery in this area. For instance, substitutions in coding regions of conserved genes have been implicated in phenotypic changes responsible for adaptation of marine mammals to an aquatic lifestyle (Foote et al., 2015; Zhou et al., 2015). Other examples of convergent phenotypes whose molecular underpinnings have been investigated include adaptations in snake and agamid lizard mitochondria (Castoe et al., 2009), echolocation in mammals (Parker et al., 2013; Thomas and Hahn, 2015; Zou and Zhang, 2015; Storz, 2016), and hemoglobin function in birds (Natarajan et al., 2016).

Several traits are also known to have convergently evolved in land plants (e.g., Li et al., 2018; Lü et al., 2018; Preite et al., 2019). One of the most notable examples is represented by the repeated evolution of the C_4_ photosynthetic pathway in flowering plants. The C_4_ pathway is a complex functional adaptation that allows for better photosynthesis efficiency under certain environmental conditions, such as dry and warm climates, high light intensity, low CO_2_ concentration, and limited availability of nutrients (Knapp and Medina, 1999; Long, 1999). The C_4_ pathway involves cytological, anatomical and metabolic modifications thought to have evolved multiple times independently in various lineages from the C_3_ type (Kellogg, 1999; Sage, 2004; Sage et al., 2011). According to phylogenetic, anatomical and biochemical evidence, the few slightly different variants of the C_4_ photosynthesis type originated more than 60 times in angiosperms (Sage et al., 2012; Heyduk et al., 2019). In grasses (family Poaceae) alone, the C_4_ pathway has evolved independently ~20 times (Grass Phylogeny Working Group II, 2012).

Transitions from C_3_ to C_4_ plants resulted from genetic changes that include nonsynonymous substitutions, gene duplications and gene expression alterations (Christin et al., 2007; Christin et al., 2013a; Christin et al., 2015; Goolsby et al., 2018; Heyduk et al., 2019). It has been suggested that the evolution of the C_4_ pathways proceeded throughout a series of evolutionary steps wherein the Kranz leaf anatomy typical of this pathway originated first, followed by changes in the expression patterns of key genes and finally by adaptive modifications of protein sequences that often result in the convergent emergence of the same amino acid replacements across C_4_ lineages (Christin et al., 2013b; Sage et al., 2012; Williams et al., 2013).

Evidence of convergent changes in proteins associated with photosynthetic processes has steadily accumulated since genomic data from multiple C_4_ lineages have become available in the past couple of decades. Most of these studies have focused on the ribulose-1,5-bisphosphate carboxylase/oxygenase (RuBisCO), a large multimeric enzyme that catalyzes the carboxylation of ribulose-1,5-bisphosphate (RuBP), allowing plants to fix atmospheric carbon (Andersson and Backlund, 2008). RuBisCO also initiates oxygenation of RuBP, which leads to a more limited production of energy and to loss of carbon in the process of photorespiration (Andersson and Backlund, 2008; Maurino and Peterhansel, 2010). RuBisCO’s limited ability to discriminate between CO_2_ and O_2_ has been attributed to the much higher CO_2_ to O_2_ atmospheric partial pressure until ~400 million years ago (Sage, 1999, 2004; Sage et al., 2012).

Previous studies have revealed multiple convergent amino acid replacements in the large RuBisCO subunit in C_4_ lineages, encoded by the chloroplast gene *rbcL* (Kapralov and Filatov, 2007; Christin et al., 2008; Kapralov et al., 2011; Kapralov et al., 2012; Piot et al., 2018). Some of these convergent replacements have been associated to positive selection of the corresponding codons in C_4_ monocot and eudicot lineages (Kapralov and Filatov, 2007; Christin et al., 2008; Kapralov et al., 2012; Piot et al., 2018). Notably, biochemical analyses have demonstrated that some recurrent amino acid changes in the large RuBisCO subunit of C_4_ plants critically alter the kinetics of RuBisCO, resulting in an accelerated rate of CO_2_ fixation at the beginning of the Calvin-Benson cycle (Studer et al., 2014). Convergent amino acid changes have also been described in enzymes that are encoded by nuclear genes and play a primary role in the C_4_ pathway, including the phosphoenolpyruvate carboxylase PEPC (Christin et al., 2007; Besnard et al., 2009), the NADP-malic enzymes NADP-me (Christin et al., 2009a), the phosphoenolpyruvate carboxykinase PEPCK (Christin et al., 2009b) and the small RuBisCO subunit (Kapralov et al., 2011).

Given the number of biochemical, physiological and anatomical traits that were affected in each evolutionary transition from C_3_ to C_4_ photosynthesis (Heyduk et al. 2019), it is likely that many genes experienced analogous selective pressures across taxa that include C_4_ plants. This could have led to the widespread occurrence of convergent amino acid replacements among a significant fraction of proteins encoded by genes involved in photosynthesis processes. A recent, important work has produced the first analysis of convergent replacements across multiple proteins involved in the metabolism of C_4_ and crassulacean acid metabolism (CAM) among species belonging to the portullugo clade (Caryophyllales). Goolsby and colleagues (2018) compared evolutionary patterns in 19 gene families with critical roles in metabolic pathways of both C_4_ and CAM plants, also known as carbon-concentration mechanisms (CCMs) genes, and in 64 non-CCM gene families. They found convergent replacements in proteins from C_4_ and CAM lineages, as well as higher levels of convergent replacements in CCM vs. non-CCM gene families (Goolsby et al., 2018). Additionally, several amino acid replacements that are prevalent among C_4_ and CAM taxa compared to C_3_ lineages were identified in this study (Goolsby et al., 2018).

Altogether, the results of this and other studies demonstrated that convergent molecular evolution occurred across multiple genes in both C_4_ and CAM groups. However, a rigorous framework to assess the full extent of molecular convergence in C_3_ to CCMs transitions has yet to be presented. For example, analyses of convergent evolution should include null hypotheses that assume no differences between taxa with and without convergence. In the case of CCMs evolution, a plausible null hypothesis consists in statistically equivalent numbers of convergent replacements between C_4_ (or CAM) lineages and C_3_ lineages.

Additionally, nonadaptive replacements should be used to normalize convergent replacements, in order to account for variation in the rates of nonsynonymous substitutions across lineages. This approach has been successfully applied in studies of molecular convergent evolution in vertebrates by assessing both convergent replacements and protein sequence changes that result in different amino acids, or *divergent replacements* (Castoe et al., 2009; Thomas and Hahn, 2015; Zou and Zhang, 2015). Furthermore, testing hypotheses about the extent of convergent molecular evolution remains particularly challenging for many nuclear genes, because of the prevalence of duplicated copies, particularly in plants (Christin et al., 2007; Goolsby et al., 2018). Single-copy nuclear or organelle genes allow to more easily recognize convergent changes and overcome possible confounding compensatory effects due to the presence of paralogous copies.

Given these premises, we sought to test if convergent amino acid changes occur more frequently in proteins encoded by chloroplast genes in a taxon that includes multiple well-characterized lineages of C_4_ and C_3_ grasses. Chloroplast proteins represent an ideal set of targets to study the role of convergent evolution in C_3_ to C_4_ transitions for a variety of reasons. First, most chloroplast proteins are involved in biochemical and biophysical processes that are critical to photosynthesis. For instance, out of ~75 functionally annotated protein-coding genes in the maize chloroplast genome, 45 genes are implicated in photosynthesis-related processes, including *rbcL*, 17 genes coding for subunits of the photosystems I and II (PS I and PS II), 12 genes coding for subunits of the NADH dehydrogenase complex, 6 genes coding for chloroplast ATPase subunits, 4 genes coding for cytochrome b6f complex subunits, and a few more genes implicated in the assembly of other protein complexes (Maier et al., 1995). Second, nonannotated orthologous copies of chloroplast genes can be readily identified across plants through sequence homology searches, taking advantage of the thousands of complete chloroplast genome sequences currently available for green plants. Third, comparative studies of convergent evolution in C_4_ photosynthesis are facilitated by detailed reconstruction of phylogenetic relationships within groups with both C_4_ and C_3_ lineages. Fourth, signatures of positive selection have been found in multiple chloroplast genes in taxa that contain both C_3_ and C_4_ plants, although only the genes *rbcL* and *psaJ*, which encodes a small subunit of the Photosystem I complex, showed evidence of adaptive changes exclusively in C_4_ lineages (Christin et al., 2008; Goolsby et al., 2018; Piot et al., 2018). Finally, most chloroplast genes occur as single copy loci, as opposed to the multiple paralogs typically present for plant genes encoded in the nucleus.

In this study, we analyzed 67 chloroplast genes from 64 grass species, including 43 C_4_ and 19 C_3_ species belonging to the PACMAD clade, named after six of its most representative subfamilies: Panicoideae, Arundinoideae, Chloridoideae, Micrairoideae, Aristidoideae and Danthonioideae. Using published phylogenetic information, we identified thirteen independent C_3_ to C_4_ transitions in this group of species. We applied a series of tests based on convergent vs. divergent amino acid replacements and determined that convergent molecular evolution occurred at a higher rate in chloroplast genes of C_4_ lineages compared to C_3_ lineages, a pattern that remained largely unchanged after excluding the RbcL protein from the convergence analyses. Our findings suggest that the evolutionary trajectories of multiple chloroplast genes have been remarkably affected during the emergence of the C_4_ adaptation in the PACMAD clade, a result that has significant implications for our understanding of C_4_ photosynthesis evolution and organelle-nucleus interactions, and for the identification of molecular changes that might be critical to the successful development of engineered C_3_ crops that incorporate carbon-concentration mechanisms.

## Methods

### Data source and filtering

We queried NCBI GenBank (Sayers et al., 2019) for complete chloroplast genome sequences of grass species that were included in phylogenetic analyses by the Grass Phylogeny Working Group II (2012) and downloaded the corresponding coding sequences. Each species was assigned to either C_3_ or C_4_ type following the results of the Grass Phylogeny Working Group II (2012). Additionally, we downloaded the coding chloroplast sequences for *Dichanthelium acuminatum, Thyridolepis xerophila, Sartidia dewinteri* and *Sartidia perrieri* (C_3_ species) (Brown and Smith, 1972; Smith and Brown, 1973; Hattersley and Stone, 1986; Hattersley et al., 1986; Besnard et al., 2014). We used the standalone blastn ver. 2.2.29+(Camacho et al., 2009) with the Expect value (E) cutoff of 1e-10 to determine putative sequence orthology with coding sequences of the *Zea mays* chloroplast genes (Maier et al., 1995). Single copy putative orthologs that were present in more than 95% of the species were retained for further analysis (Table S1).

### Multiple sequence alignment

We aligned the individual sequences using TranslatorX ver. 1.1 (Abascal et al., 2010), and further adjusted the alignments manually using BioEdit ver. 7.0.9.0 (Hall, 1999). Stop codons and sites that could not be aligned unambiguously were removed.

### Phylogeny reconstruction

We concatenated the individual sequence alignments and extracted third codon position sites for phylogeny reconstruction. We ran PartitionFinder ver. 1.1.1 (Lanfear et al., 2012) to identify the best partitioning scheme (partitioning by gene) for the downstream analysis using both Akaike information criterion (AIC) (Akaike, 1973) and Bayesian information criterion (BIC) (Schwarz, 1978). We then used maximum likelihood framework as implemented in RAxML ver. 8.2.10 (Stamatakis, 2014) to reconstruct the phylogeny. Branch support was estimated using 1,000 bootstrap replicates. *Oryza sativa* and *Brachypodium distachyon* from the BOP (Bambusoideae, Oryzoideae and Pooideae) clade were used as outgroup, whereas all ingroup species belonged to the PACMAD clade. We used FigTree ver. 1.4.0 (Rambaut, 2012) to rearrange and visualize the phylogeny, and the figures were edited further to improve readability and to indicate C_4_/C_3_ classification.

### Ancestral state reconstruction

We reconstructed ancestral states at each phylogenetic node for each individual gene using the program codeml from the software package PAML ver. 4.9a (Yang, 2007) and the basic codon substitution model (model = 0, NSsites = 0).

### Inference of convergent and divergent replacements

We extracted the reconstructed ancestral states from the codeml output. The corresponding amino acid sequences were then compared to investigate individual site changes along selected branches in the reconstructed phylogenetic tree in the context of emergence of the C_4_ trait. For each group of species descendant from a single C_4_ ancestor, we chose the branch between the most recent C_3_ ancestor and the most ancestral C_4_ node, i.e., the branch along which the C_4_ adaptation presumably emerged (referred to as “C_4_ ancestral branch” throughout this article, see Figs. 1 and 2). For C_3_ species, we chose the most ancestral branch that did not share ancestry with any C_4_ lineage (“C_3_ ancestral branch”, see Figs. 1 and 2). In either case, if only a single species was available in a given lineage, that terminal branch was used. The outgroup species (*O. sativa* and *B. distachyon*) were not included in this analysis (Fig. 1).

**Figure 1.**
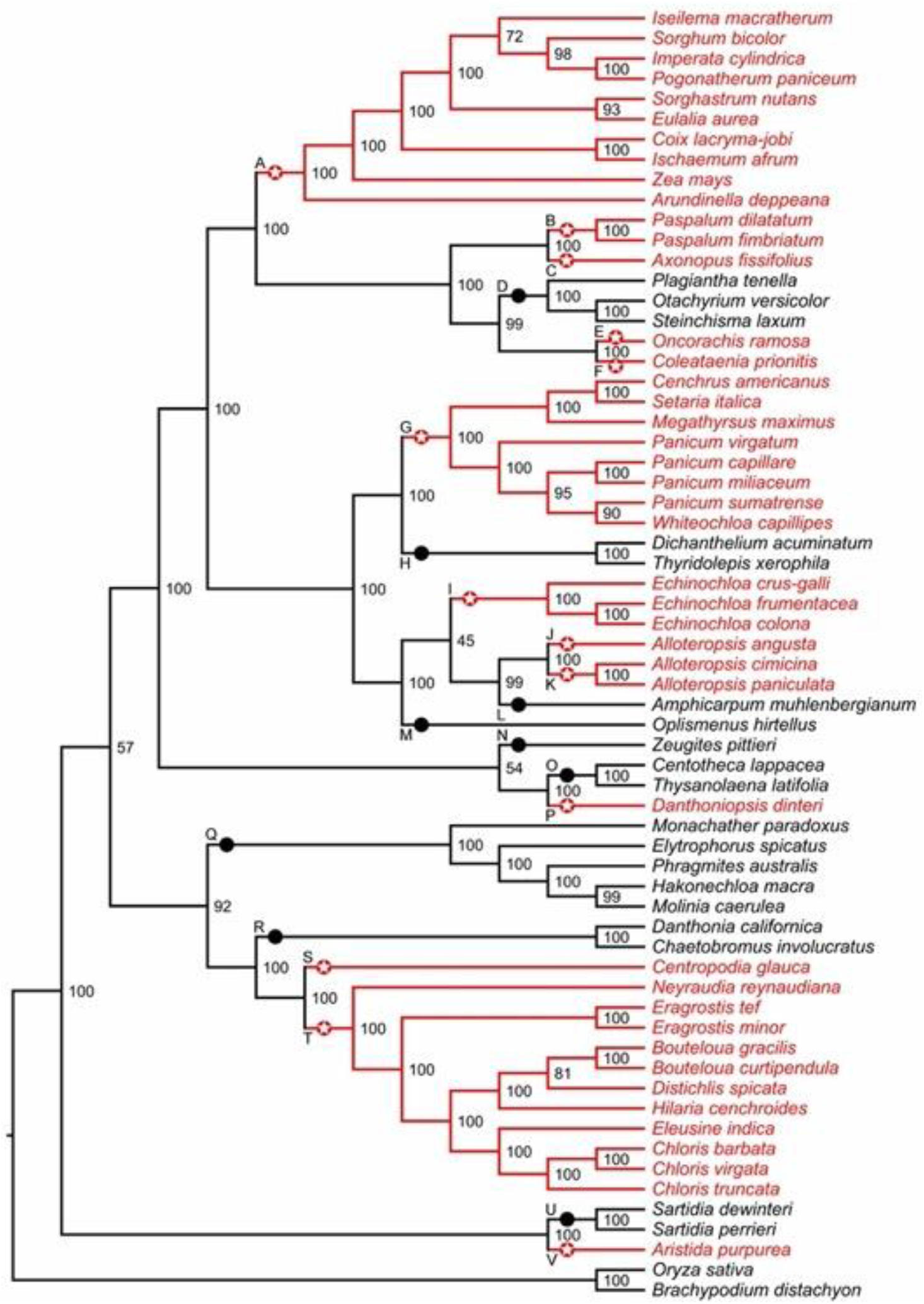
Phylogenetic relationships among 64 C_4_ and C_3_ grass species. The phylogeny tree was obtained using RAxML (GTR+Γ model) based on the third codon position sites in 67 chloroplast genes. The partitioning scheme was selected according to Akaike information criterion (AIC). C_4_ and C_3_ ancestral branches are shown in red and black, respectively. Red stars and black circles (labels A-V) indicate C_4_ and C_3_ ancestral branches, respectively. Numbers represent bootstrap support.

**Figure 2.**
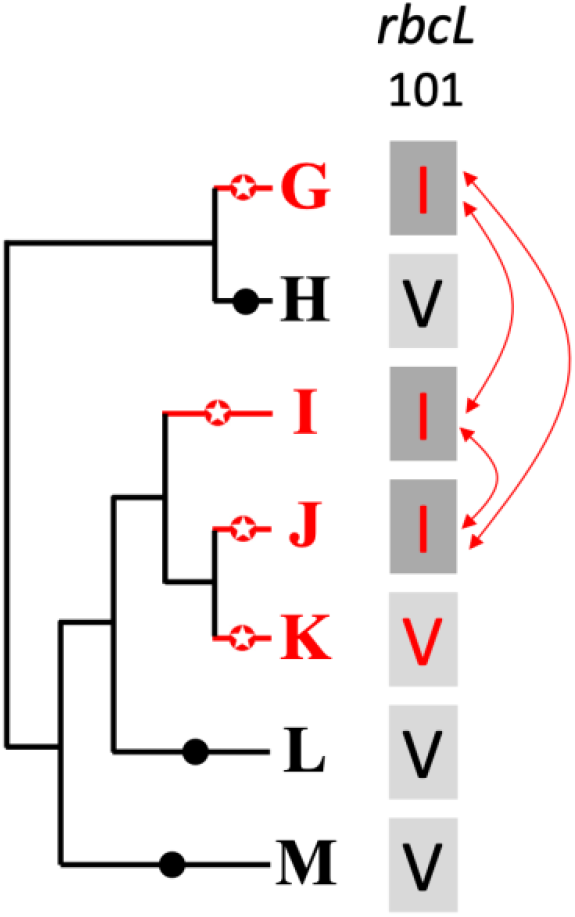
Example of ancestral C_4_ and C_3_ ancestral branches and convergent changes in C_4_ ancestral branches. Pairwise comparisons were carried out between the ancestral branches (in this example, G to M; see also Fig. 1). The C_4_ branches G, I and J shared a convergent V → I amino acid replacement (red arrows) at position 101 of the large RuBisCO subunit encoded by the gene *rbcL*. The ancestral amino acid and the convergently changed amino acidic are highlighted in light and dark gray, respectively. C_4_ and C_3_ ancestral branches are shown in red and black, respectively.

We searched for amino acid changes that occurred along pairs of ancestral branches. Replacements in both branches that resulted in the same state at a given site in the two descendants were considered convergent, regardless of whether the corresponding ancestral states of ancestral were the same or different (Castoe et al., 2009). Likewise, two replacements were considered divergent if states at the descendant orthologous sites were different, regardless of the corresponding ancestral states (Castoe et al., 2009). Although two orthologous sites, by definition, descend from one ancestral site, the actual state transitions, as well as their number, between ancestral and descendant states along a given branch are not known because the states are reconstructed only at discrete time steps (i.e., at selected nodes) and represent only those specific evolutionary time stamps. Therefore, potential intermediate stages, including a transient convergent phase, would remain undetected.

We identified putative convergent and divergent amino acid changes in each gene product individually. We summarized those data within each of the three categories: (1) two C_4_ ancestral branches (C_4_-C_4_), (2) C_3_ ancestral branch and C_4_ ancestral branch (C_3_-C_4_), and (3) two C_3_ ancestral branches (C_3_-C_3_). the Boschloo’s statistical exact unconditional test (Boschloo, 1970) was performed was performed to test the significance of the convergent replacement excess when comparing two of the three photosynthesis type pairs using the SciPy library ver. 1.7.1 in python3 (Virtanen et al. 2020).

## Data availability

Raw data, including alignments, fasta sequences, and phylogenetic analyses data, are available through the following Figshare repository: https://figshare.com/articles/dataset/Convergence-chlorplast-genes-C4-Casola-Li-2021/15180690.

## Results

### Phylogeny reconstructions

We examined 63 grass chloroplast genomes to identify gene orthologs for *Zea mays* chloroplast genes and extracted the corresponding coding and protein sequences. The resulting dataset included up to 67 DNA/protein sequences in 64 grass species that were retained for further analysis (Table S1). One to four sequences were absent in thirteen species. Out of 64 species, 43 were classified as C_4_ and 21 (including two outroup species) as C_3_. The reconstructed phylogeny is well supported, except for three branches with low to moderate bootstrap values, and it is consistent for both AIC and BIC (Fig. 1 and Figs. S1-S3). We identified thirteen C_4_ ancestral branches that represent putative C_3_ to C_4_ transitions, and nine C_3_ ancestral branches (Fig. 1).

### Convergent and divergent amino acid replacements across chloroplast proteins

We assessed the level of molecular convergence in C_3_ to C_4_ transitions by quantifying convergent and divergent amino acid replacements across the PACMAD phylogeny by performing pairwise comparisons of reconstructed ancestral sequences (Figs. 2 and 3, Table S2; see Methods).

**Figure 3.**
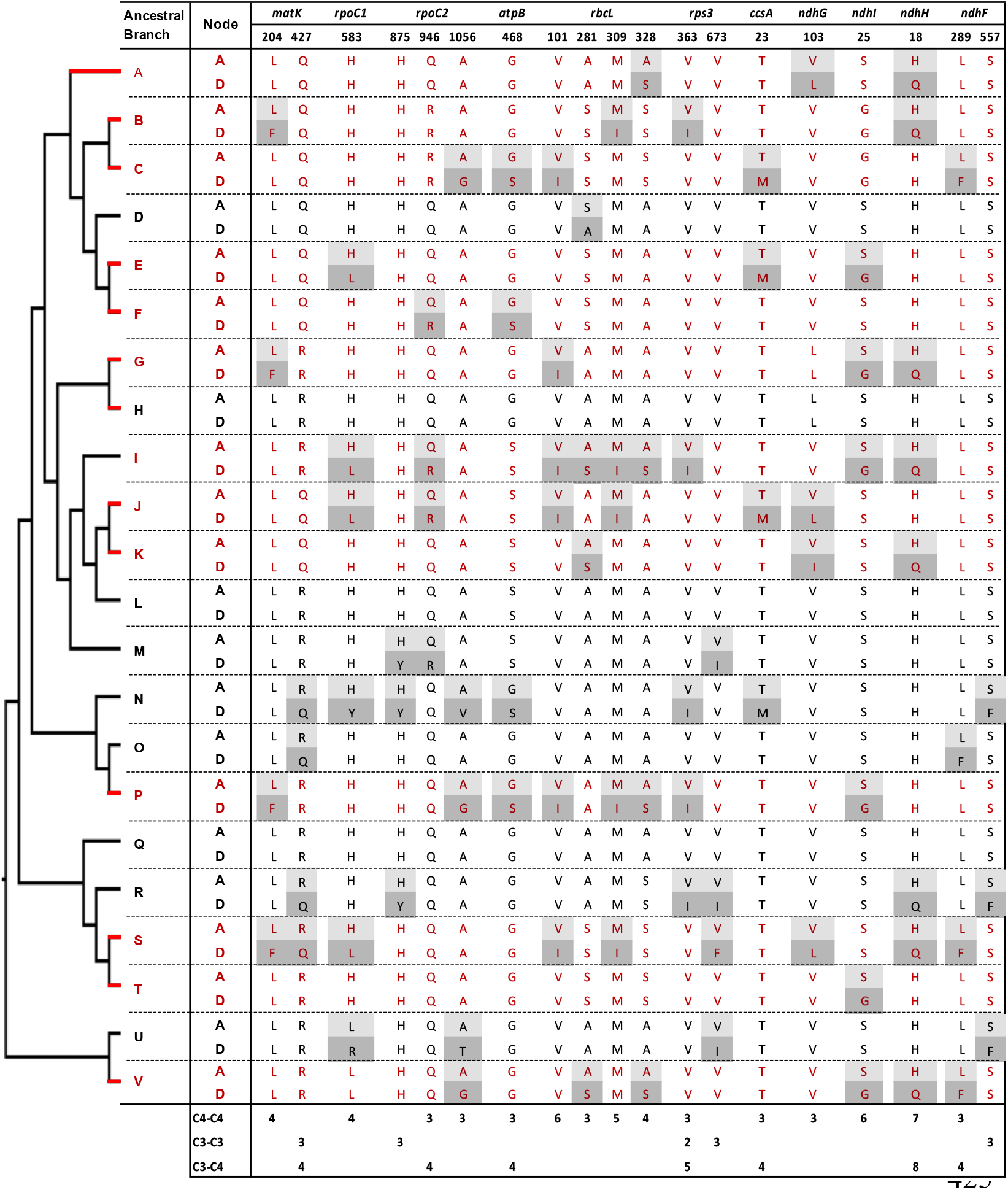
Amino acid replacements shared by at least three C_4_ or C_3_ reference branches. Ancestral (A) and derived (D) amino acids at replacement sites are shown. Site numbers correspond to the *Zea mays* orthologous sequence annotation. Red and black letters and branches represent C_4_ and C_3_ ancestral branches, respectively (see also Figs. 1 and 2).

A total of 217 sites showed at least one convergent replacement: 104 in C_4_-C_4_, 120 in C_3_-C_4_ and 34 in C_3_-C_3_ pairs. A further 201 sites exhibited one or more divergent replacements: 96 in C_4_-C_4_, 121 in C_3_-C_4_, and 39 in C_3_-C_3_ pairs (Table 1). The difference in convergent-divergent site distributions between the three photosynthesis types was not statistically significant (*P* ≥ 0.05, Boschloo’s test; Table 1).

**Table 1.** Numbers of amino acid sites and genes with convergent and divergent replacements in ancestral branch comparisons.

|  | C4-C4 |  |  | C3-C4 |  |  | C3-C3 |  |  |
| --- | --- | --- | --- | --- | --- | --- | --- | --- | --- |
|  | Con | Div | Ratio | Con | Div | Ratio | Con | Div | Ratio |
| Sites | 104 | 96 | 1.08 | 120 | 121 | 0.99 | 34 | 39 | 0.87 |
| Sites* | 80 | 64 | 1.25 | 82 | 69 | 1.19 | 17 | 16 | 1.06 |
| Genes | 24 | 23 | 1.04 | 26 | 32 | 0.81 | 13 | 17 | 0.76 |
| Genes* | 24 | 20 | 1.2 | 25 | 29 | 0.86 | 9 | 10 | 0.9 |

Comparisons were made between pairs of C4-C4, C3-C3 and C3-C4 branches. Numbers of replacements unique to a given category (\*), and the corresponding ratios *Con:Div (Ratio)*. Differences between the C3-C3 and C4-C4 categories are not statistically significant ( $P \geq 0.05$ , Boschloo's test). Con: convergent. Div: divergent.

Among the C_4_ ancestral branches, several individual sites showed high contrast in the number of branches involved in convergent and divergent replacements (Fig. 3, Tables S2 and S3). For example, seven C_4_ branches (54%) shared the H18Q replacement in the product of *ndhH*, with no divergent replacements. Six, five, and four C_4_ branches (46%, 38%, and 31%) Showed convergent replacements at three sites in the RbcL protein (V101I, M309I, and A328S, respectively). Furthermore, six C_4_ branches shared the S25G replacement in the product of *ndhI* and four L204F changes in the protein encoded by *matK*. In all these cases, there were no other convergent or divergent replacements in C_3_-C_3_ or C_3_-C_4_ branch comparisons, except for one H18Q change in NdhH in a C_3_-C_3_ branch. Two sites with convergent replacements in the proteins encoded by *ndhF* (L557F) and *rpoC2* (H875Y) were found uniquely in C_3_-C_3_ pairs, and only one site in the protein Rps3 showed convergence independently in C_4_-C_4_ and C_3_-C_3_ pairs (Fig. 3).

We then searched for convergent replacements that occurred along more than two C_4_ branches at sites that remained otherwise conserved in C_3_ and C_4_ lineages, arguing that such changes could result from selective pressure rather than drift. We identified fourteen C_4_-specific convergent sites in proteins from 8 genes: *atpB, ccsA, matK, ndhF, ndhH, ndhI, rbcL* and *rpoC2* (Table S3). Five of these sites were found in RbcL, whereas two sites were identified in both NdhF and NdhI.

### Molecular convergence in individual chloroplast proteins

Convergent and divergent amino acid replacements were detected in the products of 45 chloroplast genes, thirteen of which had at least one site with four or more replacements (Fig. 4, Table 1 and Table S2). Twenty-four genes had convergent changes in C_4_-C_4_, 26 in C_3_-C_4_, and 13 in C_3_-C_3_ types of pairs (Table 1). Although the convergent/divergent replacement ratio was higher in C_4_-C_4_ pairs than C_3_-C_4_ and C_3_-C_3_ pairs, the differences between the three photosynthesis types was not statistically significant (*P* ≥ 0.05, Boschloo’s test; Table 1). The lack of replacements was the single most common state for chloroplast proteins across photosynthesis types; however, in C_4_-C_4_ there were more genes with a higher number convergent vs. divergent replacements (Fig. 4 and Table S4).

**Figure 4.**
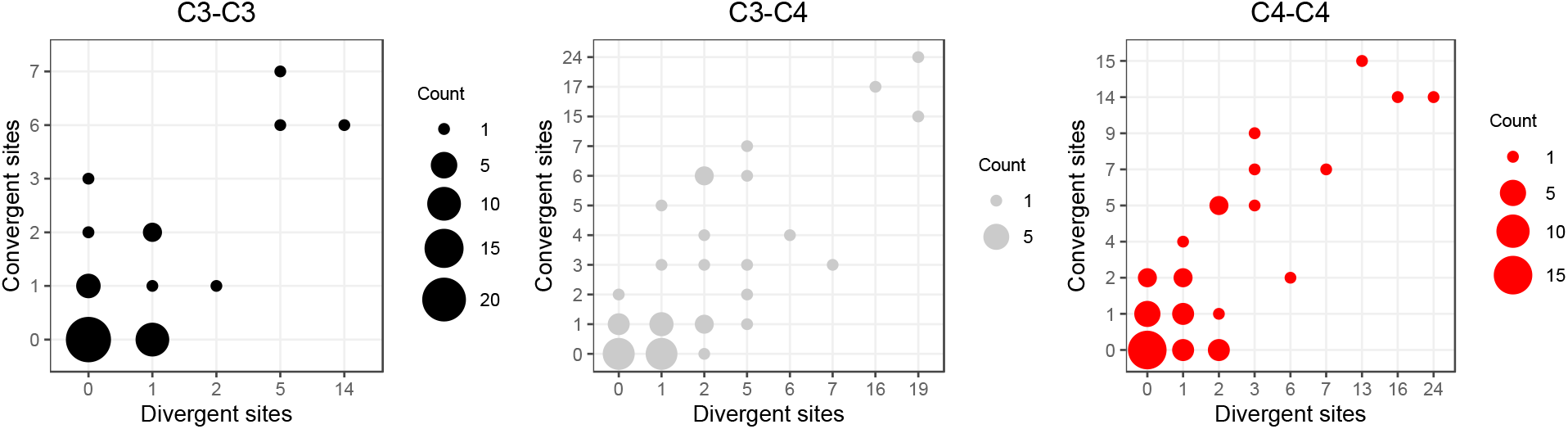
Distribution of convergent and divergent amino acid replacements in pairs of ancestral branches. Overall, 26 proteins showed a higher number of convergent vs. replacement sites, of which 16, 13 and 10 were found in C_4_-C_4_, C_3_-C_4_ and C_3_-C_3_ pairs, respectively (Fig. 5 and Table S4). We found statistically significant differences in the number of replacements between C_4_-C_4_ and C_3_-C_4_ pairs, but not C_3_-C_3_ pairs, in the products of the genes *rbcL, rpoC1* and *rpoC2* (*P* < 0.05, Boschloo’s test; Table S4). In RbcL and RpoC1, C_4_-C_4_ pairs shared much higher number of convergent replacements, whereas the opposite was true in RpoC2. RpoC1 was also the only protein showing more convergent than divergent replacements in C_4_-C_4_ pairs compared to C_3_-C_3_ and C_3_-C_4_ pairs. In C_4_-C_4_ pairs, RpoC1 shared 4 convergent and 1 divergent replacement, compared to 1 and 2 in C_3_-C_3_ pairs and 1 and 5 in C_3_-C_4_ pairs, respectively. Additionally, the proteins NdhG, NdhI, PsaI, RpoA, Rps4 and Rps11 exhibited convergent replacements only in C_4_-C_4_ pairs (Table S4). When considering the number of affected sites rather than the number of replacements, no genes showed a significantly different pattern between photosynthesis types (*P* ≥ 0.05, Boschloo’s test; Table S4).

**Figure 5.**
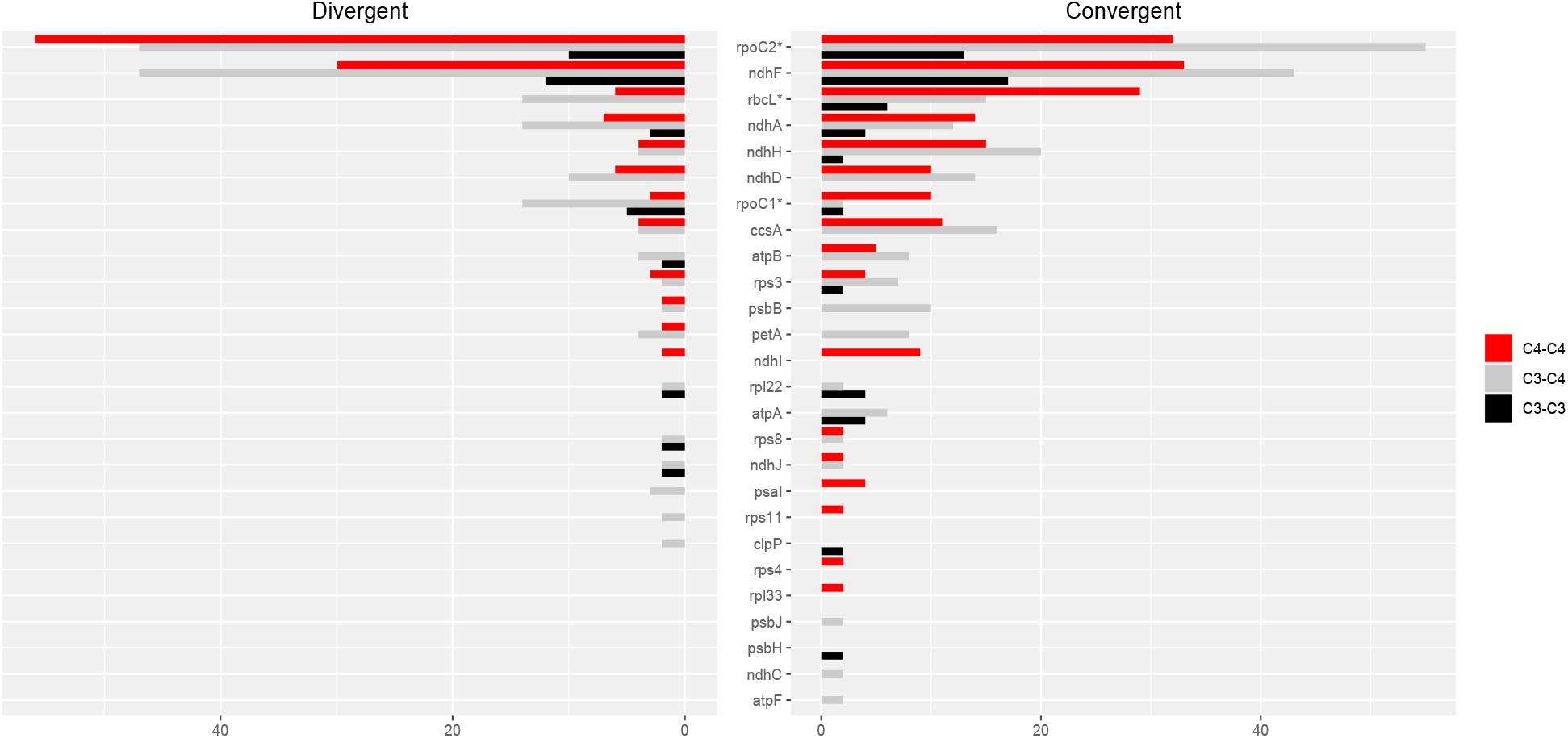
Amino acid replacements in chloroplast proteins with more convergent than divergent changes in at least one photosynthesis type. Twenty-six chloroplast proteins with more convergent than divergent changes in C_4_-C_4_, C_3_-C_4_ and/or C_3_-C_3_ pairs. Asterisks indicate proteins with significantly different replacements between C_4_-C_4_ and C_3_-C_4_ pairs.

The proteins encoded by *matK, rpoC2* and *ndhF* shared much higher numbers of both convergent and divergent replacements than other chloroplast proteins across all photosynthesis type comparisons (Table S4). Both *matK* and *ndhF* are known to be rapidly evolving and have been consistently used in low taxonomic level phylogenetic studies in flowering plants (Barthet and Hilu, 2008; Patterson and Givnish, 2002). The gene *rpoC2* has also been recently described as a useful phylogenetic marker in angiosperms (Walker et al., 2019).

### Molecular convergence across ancestral branches

The comparison of ancestral branch pairs with convergent and divergent replacements revealed remarkable differences between photosynthesis types. Overall, C_4_-C_4_ pairs of ancestral branches showed a distribution skewed toward more convergent and divergent replacements than the two other categories (Fig. 6). There were significantly fewer pairs of C_4_-C_4_ ancestral branches with no replacements and with no convergent replacements than C_3_-C_4_ and C_3_-C_3_ pairs (*P* < 0.05, Boschloo’s test; Table 2). Conversely, significantly more C_4_-C_4_ pairs shared more convergent than divergent replacements, and at least two convergent changes compared to C_3_-C_4_ and C_3_-C_3_ pairs (*P* < 0.05, Boschloo’s test; Table 2).

**Table 2.** Number of ancestral branches with convergent and divergent replacements.

|  | C<sub>4</sub>-C<sub>4</sub> | C<sub>3</sub>-C<sub>4</sub> | C<sub>3</sub>-C<sub>3</sub> |
| --- | --- | --- | --- |
| No replacements | 6 (.08) | 30 (.26) | 12 (.33) |
| No Con | 12 (.15) | 48 (.41) | 16 (.44) |
| w/Con | 66 (.85) | 69 (.59) | 20 (.56) |
| w/Div | 63 (.81) | 67 (.57) | 18 (.50) |
| Con>Div | 40 (.51) | 36 (.31) | 10 (.28) |
| Con>1 | 49 (.63) | 39 (.33) | 8 (.22) |

Comparisons were made between pairs of C<sub>4</sub>-C<sub>4</sub>, C<sub>3</sub>-C<sub>3</sub> and C<sub>3</sub>-C<sub>4</sub> branches. Proportions of pairs of ancestral branches over all branches by category are shown in parenthesis. The total number of pairs of ancestral branches are 78, 36 and 117 for C<sub>4</sub>-C<sub>4</sub>, C<sub>3</sub>-C<sub>3</sub> and C<sub>3</sub>-C<sub>4</sub> comparisons, respectively. All comparisons between C<sub>4</sub>-C<sub>4</sub> pairs and both C<sub>3</sub>-C<sub>3</sub> and C<sub>3</sub>-C<sub>4</sub> pairs were statistically significantly different ( $P < 0.05$ , Boschloo's test). No comparison between C<sub>3</sub>-C<sub>3</sub> and C<sub>3</sub>-C<sub>4</sub> pairs was statistically significant ( $P \geq 0.05$ , Boschloo's test). Con: convergent. Div: divergent. Con>Div: pairs of branches with more convergent than divergent replacements. Con>1: pairs of branches with more than one convergent replacement.

**Figure 6.**
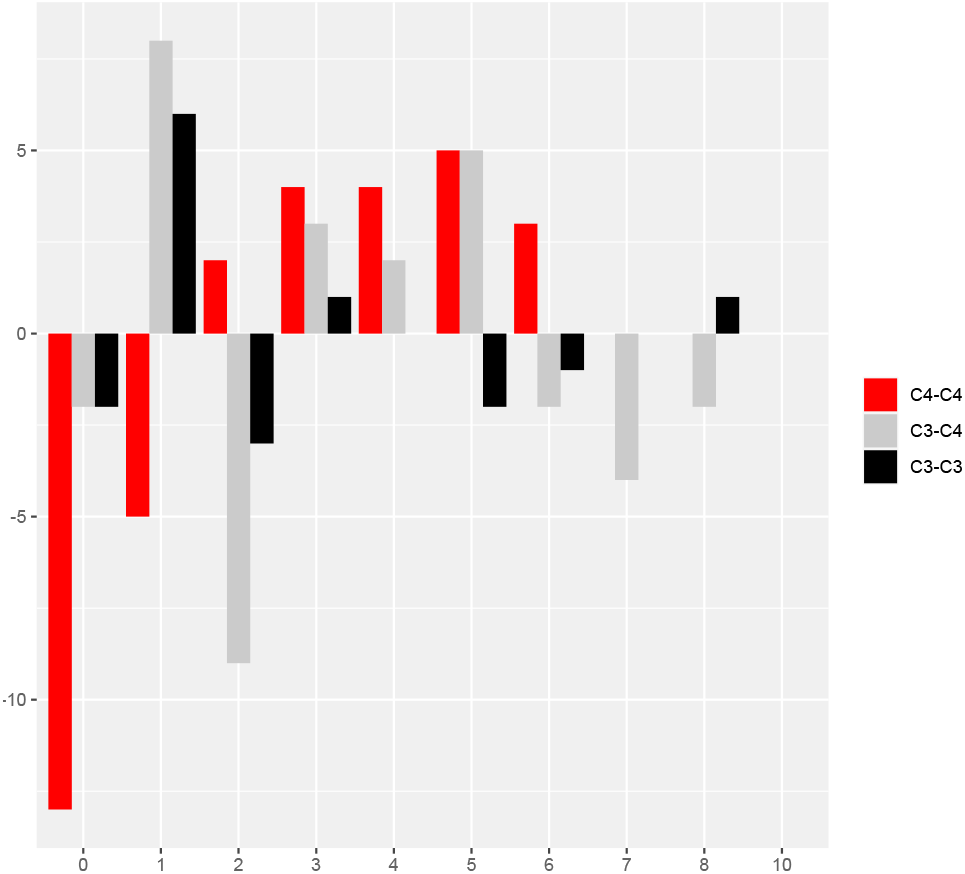
Pairs of ancestral branches by convergent and divergent replacements. Difference in the number of pairs of ancestral branches for convergent and divergent categories (0-8 and 10 replacements).

No significant difference was observed between pairs of C_3_-C_4_ and pairs of C_3_-C_3_. We found identical patterns when the same analyses were performed after excluding all replacements in the RbcL protein, except for the lack of a significant difference between C_4_-C_4_ and C_3_-C_3_ in the proportion of pairs with divergent replacements and pairs with more convergent than divergent changes (Table S6).

### Distribution of amino acid replacements across PACMAD lineages

Convergent and divergent replacements were preferentially found in specific pairs of ancestral branches. In C_4_ pairs, convergent sites were most abundant between *Danthoniopsis dinteri* and *Aristida purpurea* (ten sites, branches P and V in Fig. 1), whereas divergent sites were most common between *Centropodia glauca* and *Aristida purpurea* (ten sites, branches S and V in Fig. 1). In pairwise C_3_ branch comparisons, most convergent sites were identified between both *Zeugites pittieri* and Danthonieae (branches N and R in Fig. 1) and Danthonieae and *Sartidia* spp. (branches R and U in Fig. 1), whereas the most divergent site-rich pair was formed by *Zeugites pittieri* and *Sartidia* spp. (eight sites, branches N and U in Fig. 1; Table S5).

### Molecular convergence in the RuBisCO large subunit

We further inspected the evolution of the RuBisCO large subunit across the PACMAD clade. A total of 4 out of 9 RbcL amino acids with convergent changes in C_4_ ancestral branches—V101I, A281S, M309I and A328S—have been identified in previous studies on PACMAD grasses (Christin et al., 2008; Piot et al., 2018) as sites that experienced adaptive evolution in C_4_ species (Table 3).

**Table 3.**
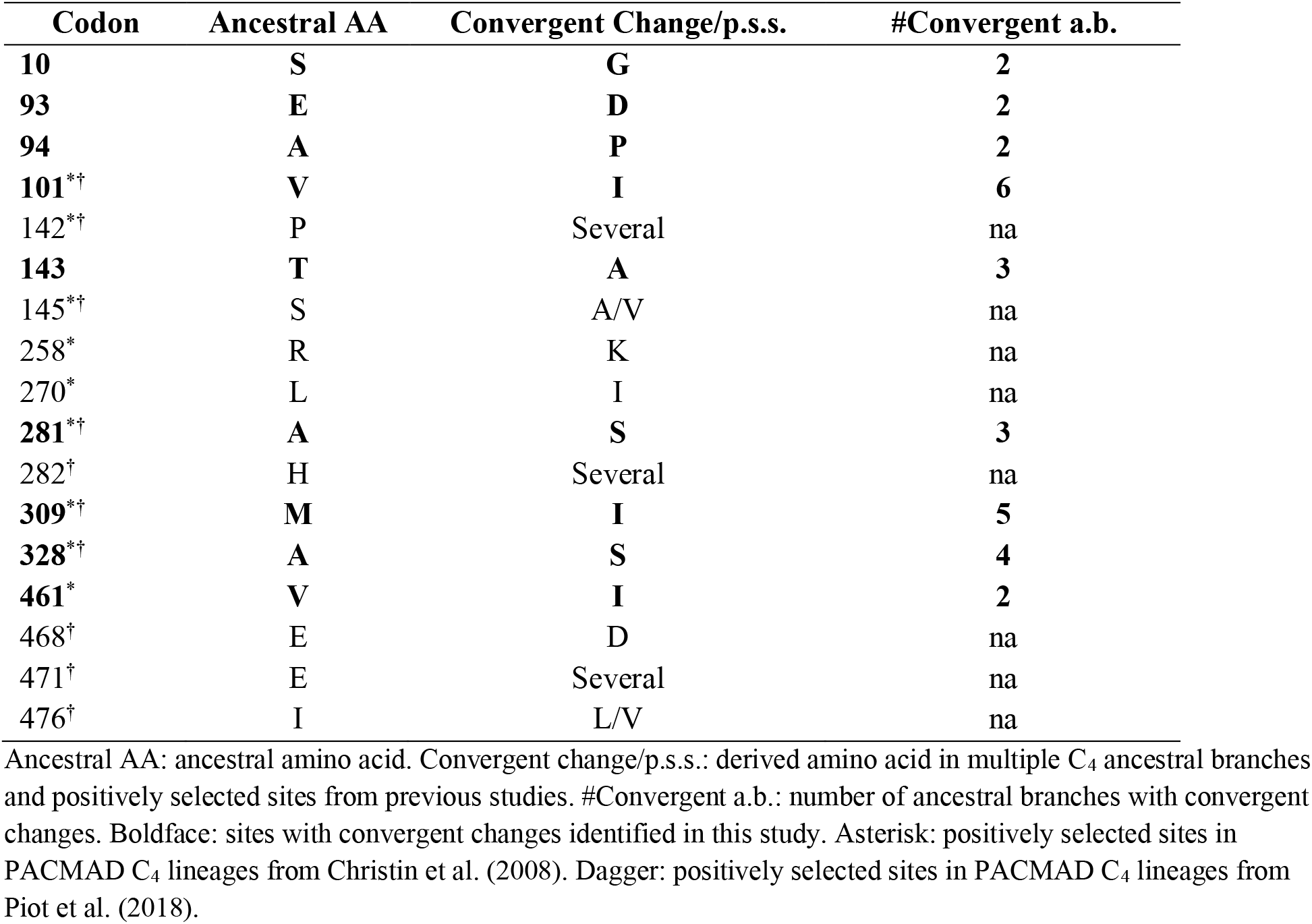
Summary of RbcL amino acid sites with signatures of convergent evolution or positive selection.

A further site, T143A, was found to evolve under positive selection in C_3_ to C_4_ transitions in monocots (Studer et al., 2014). Interestingly, an adaptive S143A replacement has also been detected in the gymnosperm *Podocarpus* (Sen et al., 2011). Three more sites with convergent replacements—at positions 93, 94 and 461—correspond to amino acids that were reported to evolve under positive selection in different groups of seed plants by Kapralov and Filatov (2007). Thus, all of the *rbcL* codons that appear to have evolved convergently among the PACMAD C_4_ lineages we have examined are also known to have experienced adaptive evolution in seed plants, but not all of them have been shown to evolve adaptively in C_4_ grasses.

## Discussion

The recurrent emergence of carbon-concentration mechanisms (CCMs) across multiple angiosperm clades in the past 35 million years represents one of the most striking examples of convergent evolution of a complex phenotypic trait. Several investigations have shown that the phenotypic parallelism across C_4_ lineages is to some extent mirrored by convergent changes in the sequence of proteins with key metabolic roles in the biochemistry of C_4_ photosynthesis, both in monocots and eudicots (Christin et al., 2007; Besnard et al., 2009, Christin et al., 2009a, Christin et al., 2009b, Kapralov et al., 2011, Goolsby et al., 2018). Furthermore, biochemical analyses have determined that some of these changes reflect adaptive shifts, as in the case of the increased availability of CO_2_ at the RuBisCO site (Studer et al., 2014). Further evidence of changes in the selective pressure associated to the C_3_ to C_4_ transitions have emerged from the detection of several positively selected sites in multiple genes associated with photosynthetic processes (Christin et al., 2008; Studer et al., 2014; Goolsby et al., 2018; Piot et al., 2018). These and other discoveries have paved the way to a more nuanced understanding of the molecular basis of phenotypic convergence in CCM plants and may accelerate the development of crop varieties with augmented resistance to high temperature and low water availability.

For these aims to be fully realized, a robust framework to assess the extent and phenotypic impact of convergent molecular changes is necessary. Along the lines of strategies applied in vertebrates research (Castoe et al., 2009, Thomas and Hahn, 2015), we presented here the results of a novel methodological approach to the study of molecular convergence in C_4_ grasses. We investigated patterns of convergent and divergent amino acid changes in nearly 70 chloroplast proteins across multiple C_4_ and C_3_ lineages in the PACMAD clade, with the goal of testing a specific hypothesis: is the evolution of chloroplast proteins showing stronger signatures of convergent amino acid replacements in C_4_ lineages compared to C_3_ lineages? This analysis also allowed us to establish if proteins other than enzymes involved in the CCM biochemistry underwent parallel amino acid changes in C_4_ lineages. Our reasoning is that many proteins expressed in the chloroplast could have experienced similar selective pressure across multiple C_3_ to C_4_ transitions and might have accumulated convergence replacements as a result.

We based our analysis on the identification of amino acid replacements shared by pairs of ancestral C_4_ branches, defined here as branches corresponding to C_3_ to C_4_ transitions in the PACMAD phylogeny. We compared these changes to those identified in ancestral C_3_ branches, namely all C_3_ lineages that include only C_3_ species (Figs. 1 and 2), and to changes found between ancestral C_3_ and C_4_ branches. For each of the three possible pairs of photosynthesis types C_4_-C_4_, C_3_-C_4_ and C_3_-C_3_, we determined the number of amino acid sites, genes and pairs of ancestral branches with convergent replacements. We detected signatures of convergent evolution in all types of datasets. First, we identified many individual replacements that emerged repeatedly and uniquely in C_4_ ancestral branches, particularly in the proteins RbcL, NdhH, NdhI and MatK. We also observed C_3_-specific convergent replacements in NdhF and RpoC2, and a case of multiple C_4_ and C_3_ convergent changes in Rps3. Additionally, we identified 8 chloroplast genes with one or more C_4_-specific convergent sites. Second, we found evidence of significantly higher rates of convergent replacements in C_4_ lineages in both RbcL and RpoC1, and several convergent replacements that occurred exclusively in C_4_-C_4_ pairs in proteins encoded by *ndhG, ndhI, psaI, rpoA, rps4* and *rps11*. These genes are involved in a variety of biological processes in the chloroplast, from the cyclic electron transport in (*ndhG* and *ndhI*) and the stabilization of (*psaI*) the photosystem I, to transcription (*rpoA* and *rpoC1*), translation (*rps4* and *rps11*) and CO_2_ fixation (*rbcL*). Third, we identified statistically significant differences in pairs of C_4_ branches with convergent replacements (Table 2). Crucially, we observed more pairs with higher convergent than divergent replacements in C_4_-C_4_ compared to both C_3_-C_3_ and C_3_-C_4,_ even after removing replacements identified in the RuBisCO large subunit, RbcL.

Altogether, these findings suggest that multiple biochemical processes occurring in the chloroplast might have experienced recurrent adaptive changes associated with the emergence of C_4_ photosynthesis. Notably, some of these proteins are not directly involved in the light-dependent or light-independent reactions of the photosynthesis, implying that processes such as the regulation of gene expression and protein synthesis in the chloroplast are also experiencing significant selective pressures during the transition from C_3_ to C_4_ plants. These results should motivate further studies to determine the prevalence of convergent amino acid replacements due transitions to CCMs among the thousands of proteins encoded by nuclear genes but expressed in the chloroplast (Jarvis and López-Juez, 2013). Although such analyses are currently hindered by the limited number of sequenced nuclear genomes in taxa with multiple C_3_ and C_4_ lineages, including the PACMAD clade, genome-wide investigations of convergent replacements will be possible in the near future given the current pace of DNA sequencing in plants.

A further important conclusion drawn from these results is that convergent replacements are not uncommon between C_3_-C_3_ and C3-C4 lineages. This is possibly due to some environmental factors affecting the evolution of chloroplast genes that are shared across grass lineages regardless of their photosynthesis type.

The analysis of individual convergent replacements in the RuBisCO large subunit both confirmed previous findings and highlighted novel potentially adaptive changes among PACMAD species. Importantly, these novel convergent replacements are known to evolve under positive selection in non-PACMAD seed plants. This underscores the potential of our approach to identify novel changes with functional significance in the transition to CCMs in grasses, as opposed to standard statistical tests of positive selection. Alternatively, some RbcL sites could experience convergence across a variety of seed plants because of selective pressure other than those associated with C_3_ to C_4_ transitions.

Overall, our results are robust to several possible confounding factors. First, we analyzed branches that are strongly supported in our phylogeny reconstruction. The phylogenetic tree built using the 67 chloroplast genes is well supported, with the exception of three branches with fairly low bootstrap support. However, all three branches are short and have minimal impact upon our conclusions regarding C_4_ evolution (Fig. 1 and Figs. S1-S3). Moreover, the tree is largely consistent with a comprehensive recent study of 250 grasses based on complete plastome data (Saarela et al., 2018). Second, by focusing only on ancestral branches and ignoring amino acid replacements that may have occurred after the divergence of species within a given C_4_ clade, our strategy provided a conservative estimate of the number of convergent changes that could have occurred during the evolution of PACMAD grasses. Third, we eliminated genes with possible paralogous copies, which could have introduced false positive replacements.

We recognize some potential caveats in our approach. By relying on a relatively small sample of PACMAD species, our statistical power to detect signatures of convergent evolution was limited. Increasing the number of ancestral C_4_ and C_3_ lineages should provide a broader representation of convergent replacements in C_4_ clades. Furthermore, we applied a strict definition of convergence that ignores changes to amino acids with similar chemical properties. We think that a conservative approach was necessary given that amino acids with similar chemical properties might have a very different functional effect on protein activity given their size and tridimensional interactions with nearby residues. Third, we assumed that all the observed convergent replacements were the result of convergent phenotypic changes, which fall under the general category of homoplasy (Avise and Robinson, 2008). However, some of these replacements could instead represent hemiplasy, or character state changes due to introgression between different C_4_ lineages, incomplete lineage sorting (ILS) of ancestral alleles or horizontal gene transfer (Avise and Robinson, 2008). Recombination between chloroplast genomes, which is required for introgression to occur, has been documented but appears to be rare (Carbonell-Caballero et al., 2015, Greiner and Sobanski, 2015, Sancho et al., 2018). Introgression or horizontal gene transfer between congeneric species has been associated to the acquisition of part of the C_4_ biochemical pathway in the PACMAD genus *Alloteropsis* (Christin et al., 2012; Olofsson et al., 2016). However, these transfers were limited to a few nuclear genes. Moreover, only a very few cases of horizontal transfer between chloroplast genomes have been reported in plants (Stegemann et al., 2012). Therefore, the contribution of hemiplasy to the observed pattern of convergent replacements in C_4_ lineages is likely to be minimal. Finally, we treated C_4_ species regardless of their photosynthesis subtype (NAPD-ME, NAD-ME and PEPCK), which is known to vary among PACMAD subfamilies (Taylor et al., 2010). We argue that our results are conservative with regard to this aspect because convergent replacements should be expected to occur more often between C_4_ groups sharing the same photosynthesis subtype.

## Conclusions

In this study, we showed that molecular convergent evolution in the form of recurrent amino acid replacements affected multiple chloroplast proteins in C_4_ lineages of the PACMAD clade of grasses. This finding significantly broadened the number of genes known to have evolved convergently in C_4_ species. We observed for the first time that genes not directly involved in photosynthesis-related processes experienced convergent changes, suggesting that future efforts should rely whenever possible on genome-wide analyses of amino acid changes rather than focus primarily on candidate key metabolic genes, similarly to previous investigations on gene expression patterns in C_4_ and CAM plants. Our methodological approach based on the comparison of convergent and divergent replacements among photosynthesis types underscores the importance of a more rigorous hypothesis-based testing of convergent evolution signatures in C_4_ plant evolution. Our results should inform more nuanced approaches to introduce CCM-like processes in C_3_ crops.

## Acknowledgements

The project was supported by the National Institute of Food and Agriculture, U.S. Department of Agriculture, under award number TEX0-1-9599, the Texas A&M AgriLife Research, and the Texas A&M Forest Service.

## Notes

https://figshare.com/articles/figure/Casola-Li_C4-convergent-chloroplast_supplementary_files/16493316

### Competing Interest Statement

The authors have declared no competing interest.

